# Nonenzymatic RNA copying with a potentially primordial genetic alphabet

**DOI:** 10.1101/2025.03.19.644114

**Authors:** Ziyuan Fang, Xiwen Jia, Yanfeng Xing, Jack W. Szostak

## Abstract

Nonenzymatic RNA copying is thought to have been responsible for the replication of genetic information during the origin of life. However, chemical copying with the canonical nucleotides (A, U, G, and C) strongly favors the incorporation of G and C and disfavors the incorporation of A and especially U, because of the stronger G:C vs. A:U base pair, and the weaker stacking interactions of U. Recent advances in prebiotic chemistry suggest that the 2-thiopyrimidines were precursors to the canonical pyrimidines, raising the possibility that they may have played an important early role in RNA copying chemistry. Furthermore, 2-thiouridine (s^2^U) and inosine (I) form by deamination of 2-thiocytidine (s^2^C) and A respectively. We used thermodynamic and crystallographic analyses to compare the I:s^2^C and A:s^2^U base pairs. We find that the I:s^2^C base pair is isomorphic and isoenergetic with the A:s^2^U base pair. The I:s^2^C base pair is weaker than a canonical G:C base pair, while the A:s^2^U base pair is stronger than the canonical A:U base pair, so that a genetic alphabet consisting of s^2^U, s^2^C, I and A generates RNA duplexes with uniform base pairing energies. Consistent with these results, kinetic analysis of nonenzymatic template-directed primer extension reactions reveals that s^2^C and s^2^U substrates bind similarly to I and A in the template, and vice versa. Our work supports the plausibility of a potentially primordial genetic alphabet consisting of s^2^U, s^2^C, I and A, and offers a potential solution to the long-standing problem of biased nucleotide incorporation during nonenzymatic template copying.

**Significance Statement:** A long-standing challenge in primordial nonenzymatic RNA copying chemistry is the biased incorporation of C and G over A and U due to differences in base pair strength. We hypothesized that 2-thiopyrimidine substitution could help overcome this bias since A:s^2^U is a stronger version of the A:U base pair, and I:s^2^C is a weaker version of the G:C base pair. This study explores the efficacy of a potentially primordial genetic alphabet consisting of s^2^U, s^2^C, A and I. Our results show that A:s^2^U and I:s^2^C pairs are isoenergetic and isomorphic. Our findings highlight the potential of this alternative genetic alphabet to yield a more balanced incorporation of all nucleotides, facilitating information propagation by nonenzymatic RNA copying during the origin of life.

## Introduction

Nonenzymatic RNA replication may have played an essential role in the transition from prebiotic chemistry to biology, before the evolution of enzymes (1). Significant progress has been made in understanding the chemistry of nonenzymatic RNA copying over the past decade. These advances include the identification of 2-aminoimidazole activated nucleotides as more effective substrates (2) and the discovery that the formation of the highly reactive 5′-5′ 2-aminoimidazolium-bridged dinucleotides (denoted by N*N) are covalent intermediates in the predominant mechanism of template copying (3, 4). Despite these advances, a key challenge persists: the biased incorporation of the canonical nucleotides during primer extension. The G:C base pair is stronger than the A:U base pair due to the presence of an additional hydrogen bond, leading to the preferential incorporation of G and C relative to A and U in nonenzymatic RNA template copying (5). In addition, the weaker stacking interactions of U make the incorporation of U in primer extension reactions particularly inefficient. We recently showed that substituting adenine with diaminopurine, which leads to a stronger base pair with U, can mitigate this bias. However, this modification falls short of achieving an even nucleotide incorporation (6); furthermore, there is as yet no plausible high yielding pathway for the prebiotic synthesis of diaminopurine nucleotides. Recently we have shown that the combined presence of random sequence oligonucleotides and activation chemistry can mitigate the nucleotide bias in RNA copying (7), presumably due to the formation of monomer-bridged-oligonucleotide substrates. While promising, this approach does suffer from the increased incorporation of mismatched nucleotides due to the ligation of mismatched oligonucleotides to the primer. We are therefore continuing to explore alternative approaches to unbiased template copying.

The 2-thiopyrimidine nucleotides may provide a distinct solution to the problem of bias in nucleotide incorporation during nonenzymatic RNA copying. Recent advances in prebiotic chemistry highlight the plausible existence of 2-thiopyrimidine nucleotides on the early Earth. The prebiotic synthesis of s^2^C has been achieved in high yield through the thiolysis of *α*-anhydro-cytidine followed by photoanomerization (8); s^2^C can then be deaminated to yield s^2^U. Similarly, inosine (I) is readily derivable by deamination of A (9). 2-thiocytidine (s^2^C) makes a weak and distorted base pair with G but can form an undistorted base pair with inosine (I) (10). The canonical pyrimidine ribonucleosides can be derived from the 2-thiopyrimidines by a variety of pathways that lead to desulfurization (8). The 2-thiopyrimidines are found in tRNA, where their presence is universally conserved across all organisms (11-14). These discoveries strongly suggest that 2-thiopyrimidines are prebiotically plausible nucleotides that could have played a significant role in the chemical evolution of life.

The influence of 2-thiouridine (s^2^U) on the thermodynamics and structure of base pairing, and on the kinetics of nonenzymatic RNA primer extension has been extensively studied (15-17). Our previous studies have demonstrated that 2-thiouridine (s^2^U) makes a stronger base pair with A (Figure 1), resulting in an increased primer extension reaction rate and better fidelity in nonenzymatic primer extension (15, 16). In contrast, less effort has been devoted to investigating 2-thiocytidine (s^2^C). The thermodynamics of C or s^2^C base pairing with either G or I has been studied in the context of the stem of an RNA stem-loop (18). In that study, the G:s^2^C base pair was weaker than the canonical G:C base pair but was almost the same as a I:s^2^C base pair, while I:C was highly destabilizing. Similarly 5-methyl-2-thiodeoxycitidine (dm^5^s^2^C) has been found to form a weaker base pair than dC with G, but a stronger base pair with I in both DNA duplexes and DNA/RNA hybrids (10). Previous work from our laboratory found that the imidazolium-bridged s^2^C substrate (s^2^C*s^2^C) has an increased maximum rate of reaction (*k*_obs max_) but a much weaker binding affinity on the -II-template compared to C*C on the -GG-template (19). The greater maximum rate of reaction may be due to s^2^C primarily adopting the 3′-endo sugar conformation (19), which is critical for nonenzymatic RNA primer extension (20).

**Figure 1.**
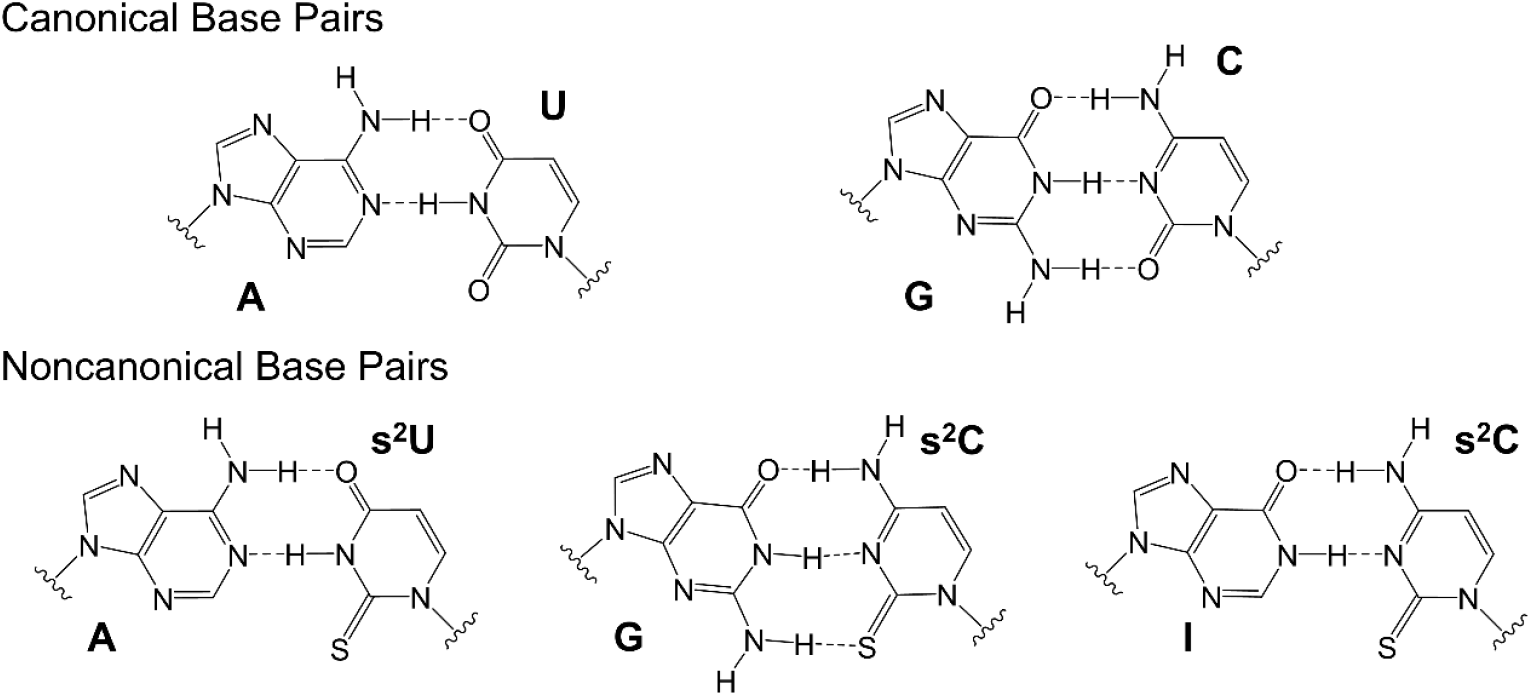
Schematic structures of the canonical A:U and G:C base pairs (top) and the noncanonical base pairs A:s^2^U, G:Cs^2^C and I:s^2^C (bottom).

Herein, we further explore the potential role of 2-thiocytidine and 2-thiouridine base pairing during nonenzymatic primer extension. We first compared the thermodynamics of base pairing of C and s^2^C to G and I, and of U and s^2^U to A. We then determined high resolution crystal structures of duplexes containing the above base pairs, all in a consistent duplex context. Finally, we examined primer extension with s^2^C substrates on G and I templates, as well as G and I substrates on C and s^2^C templates, and compare those results with s^2^U and A substrates on A and s^2^U templates. Our kinetic studies of nonenzymatic RNA primer extension reactions are consistent with the trends observed in our thermodynamic studies, and suggest that the I:s^2^C base pair, when combined with A:s^2^U, may lead to a more even incorporation of nucleobases during nonenzymatic RNA copying. We suggest that a primordial genetic alphabet consisting of A, I, s^2^U and s^2^C could potentially resolve the biased incorporation problem in nonenzymatic RNA template copying.

## Results

### Thermodynamics of Base Pairing

In order to evaluate the energetics of 2-thiopyrimidine containing base pairs and compare them with the canonical base pairs in the same context, we measured the melting temperatures (*T*_m_) of a 9-bp RNA duplex containing a variable central base pair flanked by constant sequences. *T*_m_ values were measured by variable temperature UV absorbance in 10 mM Tris-HCl at pH 8.0, 1 M NaCl, and 2.5 mM EDTA, at a series of concentrations ranging from 1.25 to 20 μM total RNA. We evaluated the thermodynamic parameters Δ*H*, Δ*S*, and Δ*G* by fitting the melting temperatures at different oligonucleotide concentrations to the Van’t Hoff equation. The resulting thermodynamic data for duplexes with six different central base pairs (G:C, G:s^2^C, I:C, I:s^2^C, A:U and A:s^2^U) are presented in Table 1. Our results are generally consistent with past research on the energetics of I:C, I:s^2^C and G:s^2^C base pairs (10, 18). Small quantitative differences in duplex stabilization by the different base pairs may reflect differences in stacking energies due to different flanking base pairs, and may also reflect differences in the denaturation of a hairpin construct (18) from our measurements on a duplex composed of two complementary strands.

**Table 1.** Thermodynamic Parameters of RNA Duplex Formation by Thermal Denaturation.

| Base pair | Sequence | $T_m^a$ (°C) | $\Delta H^b$ (kcal mol<sup>-1</sup>) | $\Delta S^b$ (kcal K<sup>-1</sup> mol<sup>-1</sup>) | $\Delta G_{25^\circ C}^c$ (kcal mol<sup>-1</sup>) |
| --- | --- | --- | --- | --- | --- |
| G:C | 5'-CUGA <u>G</u> GUAG-3'<br>3'-GACU <u>C</u> CAUC-5' | 55.6(2) | -76.2(1.3) | -0.205(4) | -15.2(3) |
| I:C | 5'-CUGA <u>I</u> GUAG-3'<br>3'-GACU <u>C</u> CAUC-5' | 46.2(2) | -74.9(1.2) | -0.208(3) | -13.0(2) |
| G:s<sup>2</sup>C | 5'-CUGA <u>G</u> GUAG-3'<br>3'-GACU <u>s<sup>2</sup>C</u> CAUC-5' | 50.1(2) | -72.8(1.0) | -0.198(3) | -13.7(2) |
| I:s<sup>2</sup>C | 5'-CUGA <u>I</u> GUAG-3'<br>3'-GACU <u>s<sup>2</sup>C</u> CAUC-5' | 52.0(2) | -79.0(1.4) | -0.216(4) | -14.6(3) |
| A:U | 5'-CUGA <u>A</u> GUAG-3'<br>3'-GACU <u>U</u> CAUC-5' | 46.8(2) | -80.9(1.0) | -0.226(3) | -13.6(2) |
| A:s<sup>2</sup>U | 5'-CUGA <u>A</u> GUAG-3'<br>3'-GACU <u>s<sup>2</sup>U</u> CAUC-5' | 53.1(2) | -77.7(1.0) | -0.211(3) | -14.7(2) |

<sup>a</sup>The reported $T_m$ was calculated from sigmoidal curves of raw thermal UV–VIS data at 5 μM total oligonucleotide, 10 mM Tris-HCl 8.0, 1 M NaCl, and 2.5 mM EDTA (Supporting Figure S1).
<sup>b</sup> $\Delta H$ and $\Delta S$ were derived from linear fits of Van't Hoff plots of $T_m^{-1}$ versus $\ln(C_T/4)$ , where $C_T$ is the total oligonucleotide concentration (Supporting Figure S2).
<sup>c</sup> $\Delta G_{25^\circ C}$ was calculated from $\Delta H$ and $\Delta S$ according to the equation $\Delta G = \Delta H - T\Delta S$ , where $T = 298.15$ K. Standard errors ( $N \geq 7$ ) are reported.

A central canonical G:C base pair confers a greater duplex stability than any other base pair. As anticipated, both the G:s^2^C and I:C base pairs exhibited lower stability compared to the canonical G:C Watson-Crick base pair. The G:s^2^C base pair leads to duplex destabilization by 1.5 kcal/mol, which appears to be the net effect of a strong enthalpic destabilization (more than 3 kcal/mol) that is partially compensated by an entropic gain. The enthalpic destabilization is likely the result of the weaker C=S···H–N hydrogen bond, due to the lower electronegativity of sulfur than oxygen, coupled with the steric distortion of the base pair caused by the larger sulfur atom (21). The lower entropic penalty for hybridization is likely due to the preorganization of s^2^C in the 3′-endo conformation (22). The non-canonical I:C base pair results in the least stable duplex, compared to all other base pairs in the center of the duplex. The duplex destabilization of 2.2 kcal/mol relative to a G:C pair is consistent with loss of the hydrogen bond between the 2-amino group of G and the 4-carbonyl of C.

The I:s^2^C base pair confers duplex stabilization that is intermediate between that of a G:C and an I:C base pair. Thus the 2-thio group of the C partially compensates for the loss of the 2-amino group of G, likely due at least in part to the lower desolvation penalty for sulfur vs. oxygen. The stability hierarchy of the six base pairs is G:C > I:s^2^C ~ A:s^2^U > G:s^2^C ~ A:U > I:C. Interestingly, the I:s^2^C and A:s^2^U base pairs contribute almost identically to duplex stabilization, with I:s^2^C being weaker than G:C while A:s^2^U is stronger than A:U. This equivalent base pair strength holds promise for achieving a more uniform product distribution in primer extension experiments.

### Structural Studies of Noncanonical Base Pairs

To further understand the structures and properties of base pairs that include s^2^C, we designed four self-complementary RNA sequences that form G:s^2^C or I:s^2^C base pairs. The sequence of the self-complementary oligonucleotide GCS1, 5′-A<u>G</u>A GAA GAU CUU CUs^2^C U-3′, assembles into a 16-mer duplex with two G:s^2^C base pairs formed from the underlined nucleotides, close to the termini of the sequence. The closely related sequence ICS1, 5′-AIA GAA GAU CUU CUs^2^C U-3′, can form two I:s^2^C base pairs in the same positions. Similarly, the sequence GCS2, 5′-AGA GAA <u>G</u>AU s^2^CUU CUC U-3′, can form two G:s^2^C base pairs from the underlined nucleotides, near the middle of the sequence, while the sequence ICS2, 5′-AGA GAA IAU s^2^CUU CUC U-3′, can form two I:s^2^C pairs at the same positions. As a reference, we also synthesized the native 16-mer self-complementary sequence and designed two sequences with A:s^2^U pairs. The sequence of the self-complementary oligonucleotide AUS1, 5′-AGA G<u>A</u>A GAU CUs^2^U CUC U-3′, assembles into a 16-mer duplex with two separated A:s^2^U base pairs formed from the underlined nucleotides. Similarly, the sequence AUS2, 5′-AGA GAA GAs^2^U CUU CUC U-3′, can form two adjacent A:s^2^U base pairs from the underlined nucleotides. All seven oligonucleotides crystallized within 2–3 days at 20 °C under their optimal crystallization conditions (Table S1), and we solved their structures by X-ray diffraction at resolutions ranging from 1.3 to 1.6 Å. Data collection and structure refinement statistics are summarized in Tables S2 and S3. We found that all seven structures adopt the same space group (*R32*). Each unit cell contains only a single RNA strand so that each duplex features two identical s^2^C or s^2^U containing base pairs.

Our crystallographic studies show that the G:s^2^C base pair has the expected Watson–Crick geometry, but slightly distorted due to the larger sulfur atom. The G:s^2^C base pairs in both GCS1 and GCS2 sequences have three hydrogen bonds and exhibit identical geometries within the resolution of our structures (Figure 2B,E). The H-bond distances between O6–N4, N1–N3, and N2–S2 in both G:s^2^C pairs are identical: 2.8, 3.0, and 3.2 Å, respectively. Compared to the N2– O2 hydrogen bond in the canonical G:C base pair (Figure 2A,C), the hydrogen bonds between N2–S2 are significantly longer, as expected due to the larger atomic radius of sulfur and the thiocarbonyl of s^2^C being a weaker hydrogen bond acceptor than the carbonyl of C. As a geometric consequence, the central N1-N3 hydrogen bond is also slightly longer in the G:s^2^C base pair. Similarly long and presumably weak hydrogen bonds involving sulfur have been seen previously in s^2^U:U and s^2^U:s^2^U pairs (17, 23). The weakened hydrogen bonds at least partly explain the thermodynamically weaker base pair between G and s^2^C.

**Figure 2.**
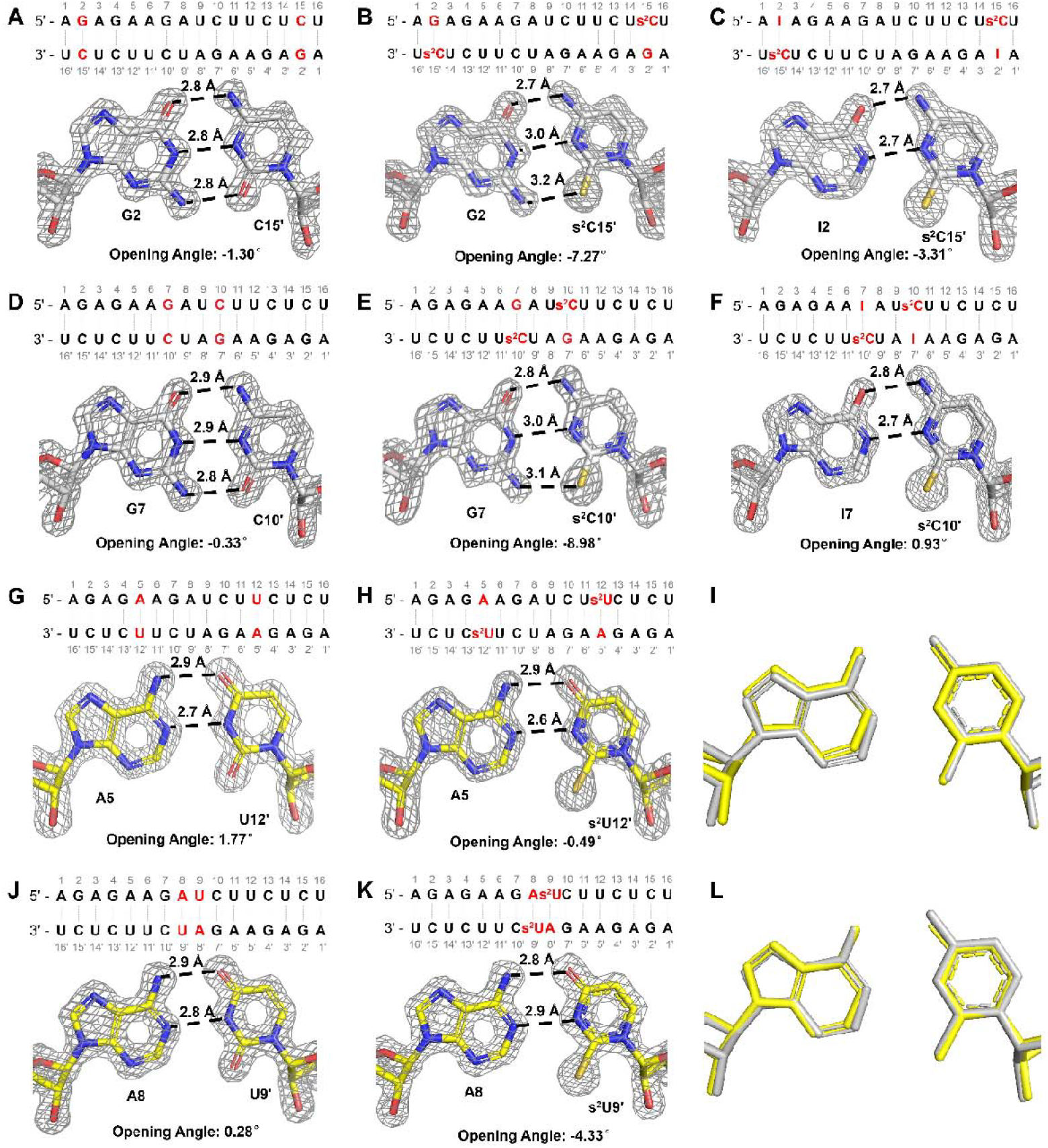
Crystal structures of G:s^2^C, I:s^2^C and A:s^2^U pairs. (A, D) Sequence, crystal structure and opening angle of the Native16 duplex containing canonical G:C pairs. (B, E) Sequence, crystal structure and opening angle of the GCS1 and GCS2 duplex containing two (B) distantly or (E) closely separated G:s^2^C pairs. (C, F) Sequence, crystal structure and opening angle of the ICS1 and ICS2 duplex containing two (C) distantly or (F) closely separated I:s^2^C pairs. (G, J) Sequence, crystal structure and opening angle of the Native16 duplex containing canonical A:U pairs. (H, K) Sequence, crystal structure and opening angle of the AUS1 and AUS2 duplex containing two (H) separated or (K) adjacent A:s^2^U pairs. (I, L) Superimposed I:s^2^C and A: s^2^U pairs in (I) GCS1 and AUS1 or (L) GCS2 and AUS2 (Silver: I:s^2^C pair; Yellow: A: s^2^U pair). Gray mesh indicates the corresponding 2F_o_–F_c_ omit maps contoured at 1.5 σ.

The I:s^2^C base pairs in the ICS1 and ICS2 sequences exhibit only two hydrogen bonds between O6–N4 and N1–N3, due to the lack of a 2-amino group on inosine (Figure 2C,F). These two hydrogen bonds are located at the same position and have similar bond lengths as in the A:s^2^U base pair (Figure 2G, J). The superimposed I:s^2^C and A:s^2^U pairs (Figure 2I, L) show that these two base pairs are isomorphic. The missing hydrogen bond is presumably the main reason that the I:s^2^C pair is weaker than the Watson-Crick G:C pair. However, the hydrogen bond between N1 on I and N3 on s^2^C has a slightly shorter length (2.7 Å) than both Watson-Crick G:C pair (2.9 Å) and G:s^2^C pair (3.0 Å). This stronger hydrogen bond may partially compensate for the loss of enthalpy due to the missing third hydrogen bond. The 2-thio group on s^2^C may increase the electron density on the aromatic ring, making N3 on s^2^C a stronger hydrogen bond acceptor than N3 on native cytidine. However, because of the geometric distortion this enhancement is not observed in the G:s^2^C base pairs.

To better understand the subtle differences between the canonical and s^2^C base pairs, we calculated the geometric parameters for all of the base pairs and base pair steps in our duplex structures (Tables S4-S13), using 3DNA (24). This analysis revealed significant changes in the opening angles of the G:s^2^C base pairs (Figure 2). Unlike the G:C pair, the G:s^2^C pair requires more space in the minor groove to accommodate the larger sulfur atom, resulting in a significantly smaller opening angle. In both the GCS1 and GCS2 sequences, the opening angles of the G:s^2^C pairs are 6-9 degrees more negative than those of canonical G:C pairs. This not only leads to a longer hydrogen bond between N2 on G and S2 on s^2^C but also imposes a geometric constraint that prevents the formation of a shorter, stronger hydrogen bond between N1 on G and N3 on s^2^C. In contrast, the absence of the 2-amino group on inosine and lack of an N2–S2 hydrogen bond results in the I:s^2^C base pair exhibiting no significant changes in the base pair opening angle, thereby making it possible to enhance the N1–N3 hydrogen bond.

Other factors may also influence the stability of RNA duplexes. For instance, sulfur is a highly polarizable atom and thus s^2^C containing base pairs could have enhanced base stacking interactions (25). However, we did not observe a significant perturbation of the overlap areas of base steps involving the s^2^C containing base pairs. Given that both G:s^2^C and I:s^2^C base pairs result in weaker hybridization than the canonical G:C base pair, changes in base stacking interactions may play a less critical role, particularly in sequences modified with a single base pair.

### Nonenzymatic Primer Extension with Thiopyrimidines, A and I

Given that the I:s^2^C and A:s^2^U base pairs appear to be isomorphic and isoenergetic, we were interested to see whether nonenzymatic template-directed primer extension with these nucleotides both as substrates and in the template would exhibit more uniform kinetics than with the canonical genetic alphabet. To address this question in a consistent context, we employed an RNA primer-helper-template duplex with a 2-nt gap in between the primer and the helper. This gap is the binding site for imidazolium-bridged dinucleotides, which are the predominant substrates for nonenzymatic primer extension with 2-aminoimidazole activated nucleotides (Figure 3A, S3) (19). To avoid any effects due to differential rates of formation or hydrolysis of imidazolium-bridged dinucleotides (abbreviated as N*N) during their formation by the spontaneous reaction of 2-aminoimidazole activated monomers (*N) with each other, we prepared and purified the bridged homo-dinucleotides G*G, I*I, C*C, and s^2^C* s^2^C as well as A*A, U*U, and s^2^U*s^2^U. We also prepared templates in which the two template nucleotides corresponding to the substrate binding site were modified with the noncanonical nucleobases. We then used these substrates and templates for primer extension reactions. We performed nonenzymatic RNA template copying reactions with complementary substrate/template pairs, as a function of substrate concentration, and fit the kinetic data using the Michaelis-Menten equation (Figure S4). The Michaelis-Menten constants (*K*_m_) and the maximum rates of reaction (*k*_obs max_) are displayed in heatmaps (Figure 3B, C).

**Figure 3.**
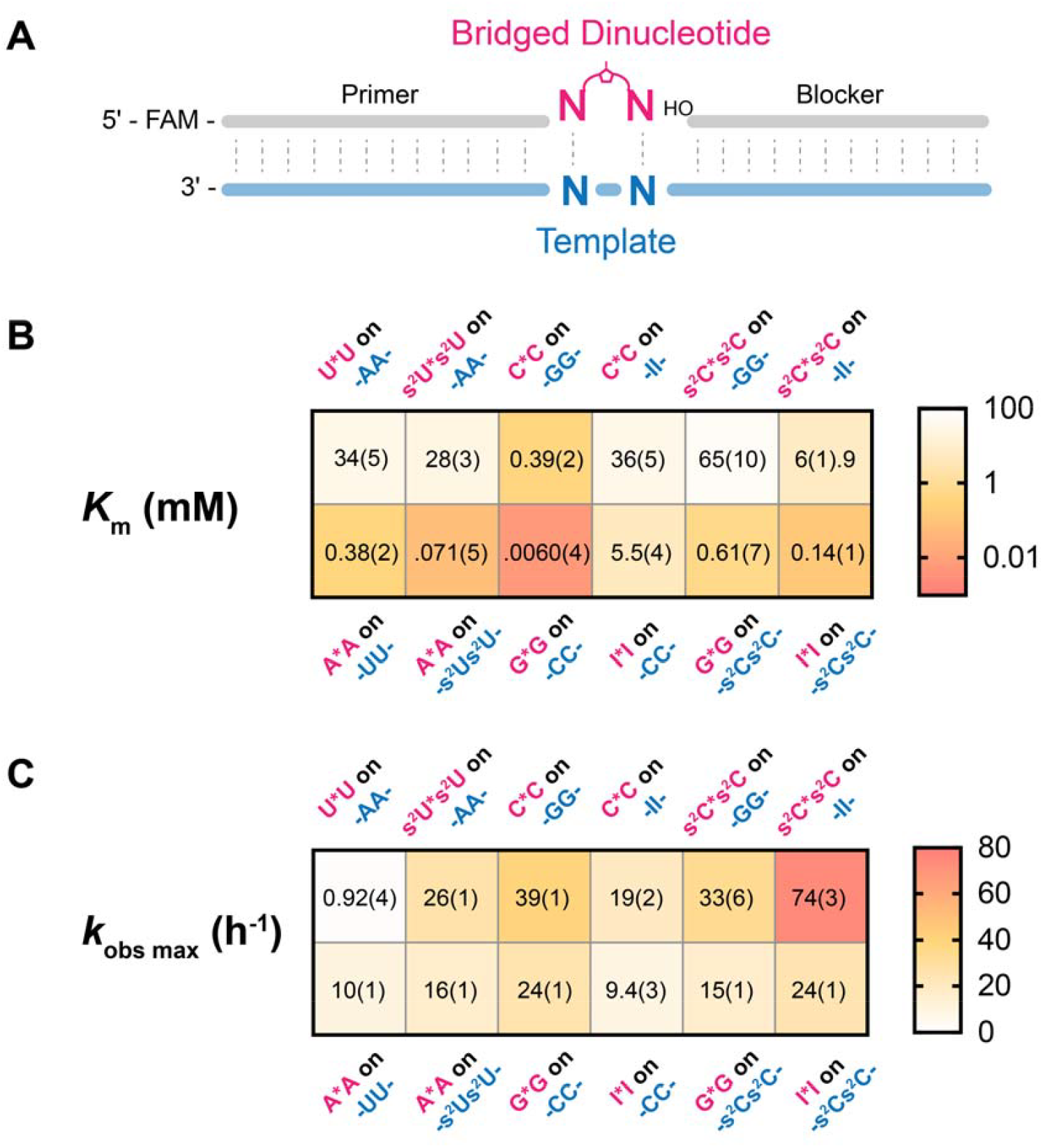
(A) Schematic representation of the nonenzymatic primer extension system. (B) Michaelis-Menten constant (*K*_m_) of the bridged dinucleotide substrates on the complementary template. (C) Observed maximum rate (*k*_obs max_) of the primer extension reactions. All reactions were performed at room temperature with 1.5 μM primer, 2.5 μM template, 3.5 μM blocker, 100 mM MgCl_2_, and 200 mM Tris-HCl pH 8.0. Standard errors N ≥ 2) are reported in parentheses.

Consistent with thermodynamic data, we observed that the binding affinities of bridged-dinucleotide substrates to the template, as reflected by their *K*_m_, followed the trend: G:C > I:s^2^C > G:s^2^C > I:C (Figure 3B). The kinetic data confirm that the weaker I:s^2^C base pair resulted in a 16~22-fold weaker binding of the s^2^C*s^2^C substrate to an II template, compared to C*C binding to a GG template. Conversely, we also see that an I*I substrate binds more weakly to a s^2^Cs^2^C template than G*G to a CC template. We were gratified to see that the binding of s^2^U*s^2^U to an AA template is only 4-5 times weaker than the binding of an s^2^C* s^2^C substrate to an II template, and that the affinity of A*A for an s^2^Us^2^U template is only two-fold stronger than the affinity of I*I for an s^2^Cs^2^C template. Notably, the maximum primer extension reaction rates for the s^2^U, A, s^2^C, I genetic alphabet system are all relatively similar. The difference between the highest reaction rate (s^2^C*s^2^C on II) and lowest reaction rate (A*A on s^2^Us^2^U) is less than 5-fold.

## Discussion

Our observation that the A:s^2^U and I:s^2^C base pairs are isoenergetic and isomorphic (Figures 2, 3 and Table 1) raises the question of whether the nucleotides A, s^2^U, I and s^2^C constitute a potentially primordial genetic alphabet. Several arguments can be made for and against the plausibility of this set of nucleotides as a progenitor of the canonical alphabet seen today in biology. On the positive side, the prebiotic synthesis of these nucleotides seems reasonable, given the current state of knowledge of prebiotic chemistry. The 2-thiopyrimidines s^2^C and s^2^U arise naturally within the cyanosulfidic reaction network as precursors of the canonical nucleotides C and U (8). In addition, s^2^U is derived from s^2^C by deamination (8), and similarly I can be derived from A by deamination (9). All three noncanonical nucleotides are found in the tRNAs of all organisms (26-28), consistent with (but not proving) the possibility that they are relics of an ancient RNA World. A second argument in favor of A, s^2^U, I and s^2^C as a primordial genetic alphabet stems from our observations of primer extension reaction kinetics. The pyrimidine substrates s^2^C and s^2^U bind with similar affinities to their complementary template sequences, as do the A and I substrates. At saturating concentrations, all four exhibit similar rates of primer extension, suggesting that this alphabet may have the capacity for relatively unbiased nonenzymatic RNA copying and replication. Further exploration of this possibility will require deep sequencing experiments under a range of conditions, to assess the extent and especially the fidelity of copying of arbitrary template sequences. Fidelity is a particular concern given the strength of the s^2^U:s^2^U self base pair, although our previous results suggest that this base pair can be outcompeted by the A:s^2^U base pair in the presence of sufficient A (23).

Despite the attractiveness of an isoenergetic base pairing landscape in terms of copying chemistry, several arguments can be made against the plausibility of a primordial genetic alphabet. First is the question of why, if life started with one alphabet, would it later switch to a different alphabet? One possibility is that nonenzymatic RNA copying may be better with the primordial alphabet, but that once a transition to ribozyme catalyzed replication had taken place, the primordial alphabet was no longer necessary. Since the canonical pyrimidines C and U are the end products of desulfurization of s^2^C and s^2^U, they might accumulate over time and be more readily available as substrates than the more transient thio-substituted intermediates. A second argument is that RNAs composed of the primordial alphabet may be less capable of forming functional folded structures such as ribozymes, due to the absence of the G:U wobble base pair (29). This possibility could be tested experimentally by in vitro selection experiments beginning with random sequence libraries composed of either the potentially primordial nucleotides or the modern nucleotides. If it is much more difficult to evolve functional ribozymes using the primordial alphabet, then life may have had to begin directly with the modern alphabet. A third argument against the primordial alphabet stems from the decreased pKa of N3 of s^2^U, which makes this nucleotide much more susceptible to alkylation by activating agents (30). However, fully plausible prebiotic activation chemistry has yet to be elucidated. The modification of s^2^U should be re-assessed as new potential activating chemistries are described.

The potentially primordial A, s^2^U, I and s^2^C genetic alphabet is likely to have strong effects on genomic replication, beyond its effects on RNA copying chemistry, due to the absence of strongly and weakly base paired regions in the genome. We have recently proposed and begun to explore a model for primordial RNA genome replication, referred to as the virtual circular genome (or VCG) model (31, 32). In this model, the genome of a protocell consists of a large collection of oligonucleotides whose sequences map to a circular consensus. The annealing of partially complementary oligonucleotides creates sites for primer extension, and repeated cycles of annealing, primer extension, and dissociation lead to lengthening of oligonucleotides and eventually genomic replication. With the canonical alphabet, regions that are AU rich will pair weakly, and thus could lead to less efficient primer extension, while regions that are GC rich could be difficult to dissociate, again leading to limited primer extension. The VCG replication strategy may therefore work better with more uniform base pair energetics. Another aspect of VCG replication that would be altered with the A, s^2^U, I and s^2^C alphabet is the generation of new primers by the template directed assembly of new short oligonucleotides (33, 34). This process is known to be more efficient with G and C than with A and U, suggesting that the canonical alphabet would lead to preferential initiation of new oligonucleotides in GC rich regions. Therefore, initiation might be more difficult in the absence of the strong GC base pairing; on the other hand, more uniform base pairing could lead to initiation at a greater number of sites in the genome. We are currently testing these aspects of VCG replication with both alphabets.

Our kinetic data for primer extension with activated homo-bridged dinucleotides reveal large differences in the affinity of bridged dinucleotides composed of purines versus pyrimidines. This trend persists even when the canonical pyrimidines are replaced with the 2-thio variants s^2^C and s^2^U, and when G is replaced by I. The very strong binding of purine dinucleotide substrates to pyrimidine template sequences may prevent the binding of pyrimidine-purine substrates to overlapping regions of the template, thereby contributing to the slow copying that is observed in mixed sequence systems. Recent advances in prebiotic chemistry have suggested the existence of potentially prebiotic pathways to the deoxypurine nucleotides (9, 35). We suggest that deoxypurine substrates may decrease the kinetic discrepancies between pyrimidine and purine substrates. Similarly, lower concentrations of ribo-purine substrates could decrease excessive template occupancy by purine substrates. However, we note that s^2^U forms a self-base pair that is energetically equivalent to a canonical A:U base pair. In the presence of equal concentrations of A, the s^2^U:s^2^U self-pair is outcompeted by the stronger A:s^2^U base pair, so that excessive misincorporation of s^2^U is avoided (23). However, if A is replaced by dA, or if the concentration of A is decreased, the level of s^2^U:s^2^U mismatch incorporation would be expected to increase. These considerations highlight the multiple trade-offs encountered during the exploration of potential scenarios for prebiotic RNA copying. We hope to gain further insight into how RNA copying may be optimized under prebiotically realistic conditions using next-generation sequencing methods (36).

## Materials and Methods

### General information

All chemicals were purchased from Sigma-Aldrich (St. Louis, MO) and used without purification unless otherwise noted. Phosphoramidites and reagents used for solid-phase RNA synthesis were purchased from ChemGenes (Wilmington, MA) and Glen Research (Sterling, MA). Preparatory-scale high performance liquid chromatography (HPLC) was carried out on an Agilent 1290 HPLC system, equipped with a preparative-scale Agilent ZORBAX Eclipse-XDB C18 column (21.2×250mm, 7 µm particle size). Purity of synthesized products was determined either by NMR or high-resolution mass spectrometry (HRMS). ^1^H and ^31^P spectra were acquired on a Bruker Ascend 9.4T/400 MHz NMR spectrometer equipped with a Bruker SampleCase Plus auto-sampler (400 MHz for ^1^H, 162 MHz for ^31^P) at 25 °C. HRMS was carried out on an Agilent 6520 QTOF LC-MS.

### Oligonucleotide synthesis

Oligonucleotides were synthesized on a K&A H-8-SE - Oligo Synthesizer, then cleaved from the solid support and deprotected with ammonium hydroxide solution at room temperature overnight. The mixtures were lyophilized, then the 2′-TBDMS protecting group was removed by treatment with triethylamine trihydrofluoride (room temperature 3 days for s^2^ C and s^2^ U-containing oligonucleotides, 65 °C 2.5 hours for canonical oligonucleotides) and purified by PAGE. The purity of oligonucleotides was confirmed by LC-MS on an Agilent 6520 TOF mass spectrometer. Oligonucleotides containing only standard nucleotides were purchased from Integrated DNA Technologies (Coralville, IA).

### Melting Temperatures of RNA Duplexes

Melting temperatures were measured using an Agilent Cary 3500 UV-Vis Spectrophotometer. For each pair of complementary oligonucleotides, samples were prepared with the desired concentration of oligonucleotide in 10 mM Tris-HCl (pH 8.0), 1 M NaCl and 2.5 mM EDTA. 200 μL mineral oil was added to the top of the RNA solution in the cuvette to prevent the evaporation of water. Melting curves were collected by following absorbance at 260 nm as a function of temperature using a temperature ramp of 0.2°C/min. The readings were collected in heating-cooling cycles with respect to a control sample containing 10 mM Tris-HCl (pH 8.0), 1 M NaCl and 2.5 mM EDTA. The melting temperatures were calculated from the interpolation of sigmoidal curves. For each concentration, two samples were prepared, and for each sample two up and down ramp cycles were carried out, generating 8 data sets for condition, i.e. four data sets from low to high temperature and four data sets from high to low temperature.

### Crystallization of RNA Duplexes

0.33 mM self-complementary 16-mer RNA sequences in nuclease-free water (Invitrogen, Waltham, MA) were heated up to 90 °C for 2 min and then slowly cooled to room temperature. Crystal Screen HT, Index HT, Natrix HT (Hampton Research, Aliso Viejo, CA) and Nuc-Pro HTS (Jena Bioscience, Jena, Germany) kits were used to screen crystallization conditions at 20 °C using the sitting-drop vapor diffusion method. An NT8 robotic system and Rock Imager (Formulatrix, Waltham, MA) were used for crystallization screening and for monitoring the crystallization process. Optimal crystallization conditions are listed in Supplementary Table S1.

### X-ray diffraction data collection, structure determination and refinement

Diffraction data were collected at a wavelength of ~ 1 Å (detailed information is listed in the SI) under a liquid nitrogen stream at 99 K on Beamline 821 or 501 at the Advanced Light Source in the Lawrence Berkeley National Laboratory (USA). The crystals were exposed for 0.25 s per image with a 0.25 Å oscillation angle. The distances between detector and the crystal were set to 180–300 mm. The data were processed by HKL2000 (37) or XDS (38). The structures were solved by molecular replacement using PHASER (39) with the structure of 3ND4 as the search model (40). All structures were refined by Phenix (41) and Refmac in CCP4i (42). After several cycles of refinement, water molecules and metal atoms with well defined density were added in Coot (43). Data collection, phasing, and refinement statistics of the determined structures are listed in Supplementary Tables S2 and S3.

Synthesis, purification and characterization of 5[-5[imidazolium bridged dinucleotides (N*N) The synthesis and purification of 2-aminoimidazolium bridged dinucleotides (A*A, U*U, G*G, C*C, I*I, s^2^C*s^2^C and s^2^U*s^2^U) were carried out as previously described (19). Characterization of these bridged dinucleotide by NMR and HRMS can be found in the supporting information.

### Nonenzymatic primer extension reactions

Annealing mixtures containing primer/template/blocker complexes were prepared at 5X final concentration: 7.5 μM primer, 12.5 μM template, 17.5 μM blocker, 50 mM Tris-Cl pH 8.0, 50 mM NaCl, and 1 mM EDTA. The solution was heated to 85°C for 30 seconds and then gradually cooled to 25°C at a rate of 0.1°C per second using a thermal cycler. This annealed mixture was then diluted 5-fold with a buffer containing 240 mM Tris-Cl pH 8.0, and 125 mM MgCl_2_ to achieve final concentrations of 1.5 μM primer, 2.5 μM template, 3.5 μM blocker, 200 mM Tris-Cl pH 8.0, and 100 mM MgCl_2_. Freshly prepared stock solutions of bridged dinucleotides at 2X desired final concentrations were added to the annealed primer/template/blocker solution to initiate templated primer extension reactions. At each time point, an 0.5 μL aliquot was added to 25 μL of quenching buffer, which contained 25 mM EDTA, 1X TBE, and 4 μM of a DNA sequence complementary to the template, in formamide. Oligonucleotide sequences are provided in Tables S14 and S15.

## Acknowledgments

J.W.S. is an Investigator of the Howard Hughes Medical Institute. This work was supported in part by grants from the NSF (2104708), the Sloan Foundation (19518) and the Moore Foundation (11479) to J.W.S. The authors thank the staff at the Advanced Light Source (ALS) beamline 821. The Berkeley Center for Structural Biology is supported in part by the Howard Hughes Medical Institute. The Advanced Light Source is a Department of Energy Office of Science User Facility under Contract No. DE-AC02-05CH11231. The ALS-ENABLE beamlines are supported in part by the National Institutes of Health, National Institute of General Medical Sciences, grant P30 GM124169.

## References

1. J. W. Szostak, The eightfold path to non-enzymatic RNA replication. Journal of Systems Chemistry 3, 1–14 (2012).

2. L. Li et al., Enhanced nonenzymatic RNA copying with 2-aminoimidazole activated nucleotides. Journal of the American Chemical Society 139, 1810–1813 (2017).

3. T. Walton, J. W. Szostak, A highly reactive imidazolium-bridged dinucleotide intermediate in nonenzymatic RNA primer extension. Journal of the American Chemical Society 138, 11996–12002 (2016).

4. W. Zhang, T. Walton, L. Li, J. W. Szostak, Crystallographic observation of nonenzymatic RNA primer extension. Elife 7, e36422 (2018).

5. D. Duzdevich et al., Competition between bridged dinucleotides and activated mononucleotides determines the error frequency of nonenzymatic RNA primer extension. Nucleic Acids Research 49, 3681–3691 (2021).

6. X. Jia et al., Diaminopurine in Nonenzymatic RNA Template Copying. Journal of the American Chemical Society (2024).

7. D. Duzdevich, C. E. Carr, B. W. Colville, H. R. Aitken, J. W. Szostak, Overcoming nucleotide bias in the nonenzymatic copying of RNA templates. Nucleic Acids Research, gkae982 (2024).

8. J. Xu et al., A prebiotically plausible synthesis of pyrimidine β-ribonucleosides and their phosphate derivatives involving photoanomerization. Nature chemistry 9, 303–309 (2017).

9. J. Xu et al., Selective prebiotic formation of RNA pyrimidine and DNA purine nucleosides. Nature 582, 60–66 (2020).

10. A. Ohkubo et al., Formation of new base pairs between inosine and 5-methyl-2-thiocytidine derivatives. Organic & Biomolecular Chemistry 10, 2008–2010 (2012).

11. G. Kawai et al., Conformational rigidity of specific pyrimidine residues in tRNA arises from posttranscriptional modifications that enhance steric interaction between the base and the 2’-hydroxyl group. Biochemistry 31, 1040–1046 (1992).

12. J. E. Jackman, J. D. Alfonzo, Transfer RNA modifications: nature’s combinatorial chemistry playground. Wiley Interdisciplinary Reviews: RNA 4, 35–48 (2013).

13. M. Helm, J. D. Alfonzo, Posttranscriptional RNA modifications: playing metabolic games in a cell’s chemical Legoland. Chemistry & biology 21, 174–185 (2014).

14. E. M. Phizicky, A. K. Hopper, tRNA biology charges to the front. Genes & development 24, 1832–1860 (2010).

15. A. T. Larsen, A. C. Fahrenbach, J. Sheng, J. Pian, J. W. Szostak, Thermodynamic insights into 2-thiouridine-enhanced RNA hybridization. Nucleic acids research 43, 7675–7687 (2015).

16. B. D. Heuberger, A. Pal, F. Del Frate, V. V. Topkar, J. W. Szostak, Replacing uridine with 2-thiouridine enhances the rate and fidelity of nonenzymatic RNA primer extension. Journal of the American Chemical Society 137, 2769–2775 (2015).

17. J. Sheng, A. Larsen, B. D. Heuberger, J. C. Blain, J. W. Szostak, Crystal structure studies of RNA duplexes containing s2U: A and s2U: U base pairs. Journal of the American Chemical Society 136, 13916–13924 (2014).

18. N. A. Siegfried, R. Kierzek, P. C. Bevilacqua, Role of unsatisfied hydrogen bond acceptors in RNA energetics and specificity. Journal of the American Chemical Society 132, 5342–5344 (2010).

19. D. Ding, L. Zhou, C. Giurgiu, J. W. Szostak, Kinetic explanations for the sequence biases observed in the nonenzymatic copying of RNA templates. Nucleic Acids Research 50, 35–45 (2022).

20. C. Giurgiu et al., Structure–Activity Relationships in Nonenzymatic Template-Directed RNA Synthesis. Angewandte Chemie International Edition 60, 22925–22932 (2021).

21. M. Sundaralingam, G. H. Lin, S. Arora, Stereochemistry of nucleic acids and their constituents. XV. Crystal and molecular structure of 2-thiocytidine dihydrate, a minor constituent of transfer ribonucleic acid. Journal of the American Chemical Society 93, 1235–1241 (1971).

22. E. T. Kool, Preorganization of DNA: design principles for improving nucleic acid recognition by synthetic oligonucleotides. Chemical reviews 97, 1473–1488 (1997).

23. D. Ding et al., Unusual Base Pair between Two 2-Thiouridines and Its Implication for Nonenzymatic RNA Copying. Journal of the American Chemical Society 146, 3861–3871 (2024).

24. G. Zheng, X.-J. Lu, W. K. Olson, Web 3DNA—a web server for the analysis, reconstruction, and visualization of three-dimensional nucleic-acid structures. Nucleic acids research 37, W240–W246 (2009).

25. S. K. Mazumdar, W. Saenger, K. H. Scheit, Molecular structure of poly-2-thiouridylic acid, a double helix with non-equivalent polynucleotide chains. Journal of Molecular Biology 85, 213–229 (1974).

26. A. Noma, Y. Sakaguchi, T. Suzuki, Mechanistic characterization of the sulfur-relay system for eukaryotic 2-thiouridine biogenesis at tRNA wobble positions. Nucleic acids research 37, 1335–1352 (2009).

27. S. Vangaveti et al., A structural basis for restricted codon recognition mediated by 2-thiocytidine in tRNA containing a wobble position inosine. Journal of molecular biology 432, 913–929 (2020).

28. F. Tuorto, F. Lyko, Genome recoding by tRNA modifications. Open biology 6, 160287 (2016).

29. S. Halder, D. Bhattacharyya, RNA structure and dynamics: a base pairing perspective. Progress in Biophysics and Molecular Biology 113, 264–283 (2013).

30. E. Sochacka et al., C5-substituents of uridines and 2-thiouridines present at the wobble position of tRNA determine the formation of their keto-enol or zwitterionic forms-a factor important for accuracy of reading of guanosine at the 3′-end of the mRNA codons. Nucleic Acids Research 45, 4825–4836 (2017).

31. L. Zhou, D. Ding, J. W. Szostak, The virtual circular genome model for primordial RNA replication. Rna 27, 1–11 (2021).

32. D. Ding, L. Zhou, S. Mittal, J. W. Szostak, Experimental tests of the virtual circular genome model for nonenzymatic RNA replication. Journal of the American Chemical Society 145, 7504–7515 (2023).

33. T. Inoue, L. E. Orgel, Substituent control of the poly (C)-directed oligomerization of guanosine 5’-phosphoroimidazolide. Journal of the American Chemical Society 103, 7666–7667 (1981).

34. T. Inoue et al., Template-directed synthesis on the pentanucleotide CpCpGpCpC. Journal of molecular biology 178, 669–676 (1984).

35. J. Xu, N. J. Green, C. Gibard, R. Krishnamurthy, J. D. Sutherland, Prebiotic phosphorylation of 2-thiouridine provides either nucleotides or DNA building blocks via photoreduction. Nature chemistry 11, 457–462 (2019).

36. D. Duzdevich, C. E. Carr, J. W. Szostak, Deep sequencing of non-enzymatic RNA primer extension. Nucleic acids research 48, e70–e70 (2020).

37. Z. Otwinowski, W. Minor, “Processing of X-ray diffraction data collected in oscillation mode” in Methods in enzymology. (Elsevier, 1997), vol. 276, pp. 307–326.

38. W. Kabsch, XDS. Acta Crystallographica Section D: Biological Crystallography 66, 125–132 (2010).

39. A. J. McCoy et al., Phaser crystallographic software. Journal of applied crystallography 40, 658–674 (2007).

40. B. H. Mooers, A. Singh, The crystal structure of an oligo (U): pre-mRNA duplex from a trypanosome RNA editing substrate. Rna 17, 1870–1883 (2011).

41. D. Liebschner et al., Macromolecular structure determination using X-rays, neutrons and electrons: recent developments in Phenix. Acta Crystallographica Section D: Structural Biology 75, 861–877 (2019).

42. G. N. Murshudov, A. A. Vagin, E. J. Dodson, Refinement of macromolecular structures by the maximum-likelihood method. Acta Crystallographica Section D: Biological Crystallography 53, 240–255 (1997).

43. P. Emsley, B. Lohkamp, W. G. Scott, K. Cowtan, Features and development of Coot. Acta Crystallographica Section D: Biological Crystallography 66, 486–501 (2010).

