## Supplementary material for "Nonenzymatic RNA copying with a potentially primordial genetic alphabet": SI

PNAS

**This PDF file includes:**

Synthesis and characterization of homo-dinucleotides
Figures S1 to S4
Tables S1 to S15

### 1.Synthesis and characterization of homo-bridged-dinucleotides

The following 5′-5′-2-aminoimidazolium-bridged-dinucleotides (A*A, C*C, G*G, U*U, I*I, $s^2C*s^2C$, $s^2U*s^2U$) were synthesized following a previously reported procedure (1). Details of the nuclear magnetic resonance (NMR) spectra and high-resolution mass spectrometry (HRMS) are provided below.

**1.1. 1,3-di-(adenosine-5'phosphoryl)-2-aminoimidazolium (A*A)**

**$^1$H NMR** (400 MHz, $D_2O$) δ 8.12 (s, 2H), 8.07 (s, 2H), 6.74 – 6.71 (m, 2H), 5.87 (d, J = 4.8 Hz, 2H), 4.59 (t, J = 4.8 Hz, 2H), 4.37 (t, J = 4.9 Hz, 2H), 4.16 – 4.06 (m, 4H), 4.05 – 3.98 (m, 2H). Peaks corresponding to residual TEAB observed at 3.20 and 1.27 ppm.

**$^{31}$P NMR** (162 MHz, $D_2O$) δ -12.81. (The purity is 100% with no detectable activated monomer peak)

**HRMS** (Q-TOF) m/z: $[M - H]^-$ Calcd. for $C_{23}H_{28}N_{13}O_{12}P_2$ 740.1456; Found: 740.1575.

**1.2.1 1,3-di-(cytidine-5'phosphoryl)-2-aminoimidazolium (C*C)**

**$^1$H NMR** (400 MHz, $D_2O$) δ 7.64 (d, J = 7.4 Hz, 2H), 6.93 (s, 2H), 6.02 (d, J = 7.5 Hz, 2H), 5.82 (d, J = 3.2 Hz, 2H), 4.25 – 4.11 (m, 10H). Peaks corresponding to residual TEAB observed at 3.20 and 1.27 ppm.

**$^{31}$P NMR** (162 MHz, $D_2O$) δ -12.82. (The purity is 100% with no detectable activated monomer peak)

**HRMS** (Q-TOF) m/z: $[M - H]^-$ Calcd. for $C_{21}H_{28}N_9O_{14}P_2$ 692.1231; Found: 692.1348.

**1.3 1,3-di-(guanosine-5'phosphoryl)-2-aminoimidazolium (G*G)**

**$^1$H NMR** (400 MHz, $D_2O$) δ 7.88 (s, 2H), 6.70 – 6.68 (m, 2H), 5.75 (d, J = 5.2 Hz, 2H), 4.65 (t, J = 5.1 Hz, 2H), 4.39 (t, J = 5.0 Hz, 2H), 4.14 – 4.07 (m, 4H), 4.03 – 3.96 (m, 2H). Peaks corresponding to residual TEAB observed at 3.19 and 1.27 ppm.

**$^{31}$P NMR** (162 MHz, $D_2O$) δ -12.84. (The purity is 100% with no detectable activated monomer peak)

**HRMS** (Q-TOF) m/z: $[M - H]^-$ Calcd. for $C_{23}H_{28}N_{13}O_{14}P_2$ 772.1354; Found: 772.1491.

**1.4. 1,3-di-(uridine-5'phosphoryl)-2-aminoimidazolium (U*U)**

**$^1$H NMR** (400 MHz, $D_2O$) δ 7.64 (d, J = 8.4 Hz, 2H), 6.95 (t, J = 2.0 Hz, 2H), 5.88 (d, J = 7.6 Hz, 2H), 5.82 (d, J = 4.3 Hz, 2H), 4.27 (t, J = 4.9 Hz, 2H), 4.21 (t, J = 5.3 Hz, 2H), 4.19 – 4.16 (m, 4H), 4.15 – 4.11 (m, 2H). Peaks corresponding to residual TEAB observed at 3.20 and 1.27 ppm.

**$^{31}$P NMR** (162 MHz, $D_2O$) δ -12.87. (The purity is 100% with no detectable activated monomer peak)

**HRMS** (Q-TOF) m/z: $[M - H]^-$ Calcd. for $C_{21}H_{26}N_7O_{16}P_2$ 694.0932; Found: 694.1042.

**1.5. 1,3-di-(inosine-5'phosphoryl)-2-aminoimidazolium (I*I)**

**$^1$H NMR** (400 MHz, $D_2O$) δ 8.15 (s, 2H), 6.71 (t, *J* = 2.0 Hz, 2H), 5.90 (d, *J* = 4.9 Hz, 2H), 4.77 (s, 2H), 4.63 (t, J = 5.2 Hz, 2H), 4.40 (t, J = 4.8 Hz, 2H), 4.08-4.16 (m, 4H), 4.07-3.98 (m, 2H). Peaks corresponding to residual TEAB observed at 3.19 and 1.27 ppm.

$^{31}$**P NMR** (162 MHz, $D_2O$) δ -12.90. (The purity is 100% with no detectable activated monomer peak)

**HRMS** (Q-TOF) m/z: [M − H]$^{-}$ Calcd. for $C_{23}H_{26}N_{11}O_{14}P_2$ 742.1141; Found: 742.1248.

**1.6. 1,3-di-(2-thiocytidine-5'phosphoryl)-2-aminoimidazolium ($s^2C$*$s^2C$)**

$^{1}$**H NMR** (400 MHz, $D_2O$) δ 7.81 (d, J = 7.5 Hz, 2H), 7.00 (s, 2H), 6.52 (s, 2H), 6.29 (d, J = 7.6 Hz, 2H), 4.39 – 4.04 (m, 10H). Peaks corresponding to residual TEAB observed at 3.20 and 1.27 ppm.

$^{31}$**P NMR** (162 MHz, $D_2O$) δ -12.86. **(**The purity is 100% with no detectable activated monomer peak)

**HRMS** (Q-TOF) m/z: [M − H]$^{-}$ Calcd. for $C_{21}H_{28}N_9O_{12}S_2P_2$ 724.0780; Found: 724.0881.

**1.7 1,3-di-(2-thiouridine-5'phosphoryl)-2-aminoimidazolium ($s^2U$*$s^2U$)**

$^{1}$**H NMR** (400 MHz, $D_2O$) δ 7.75 (d, J = 8.3 Hz, 2H), 7.00 (s, 2H), 6.56 (s, 2H), 6.10 (d, J = 8.6 Hz, 2H), 4.30 – 4.18 (m, 6H), 4.18 – 4.12 (m, 4H). Peaks corresponding to residual TEAB observed at 3.19 and 1.27 ppm.

$^{31}$**P NMR** (162 MHz, $D_2O$) δ -12.94. (The purity is 100% with no detectable activated monomer peak)

**HRMS** (Q-TOF) m/z: [M − H]$^{-}$ Calcd. for $C_{21}H_{26}N_7O_{14}S_2P_2$ 726.0460; Found: 726.0574.

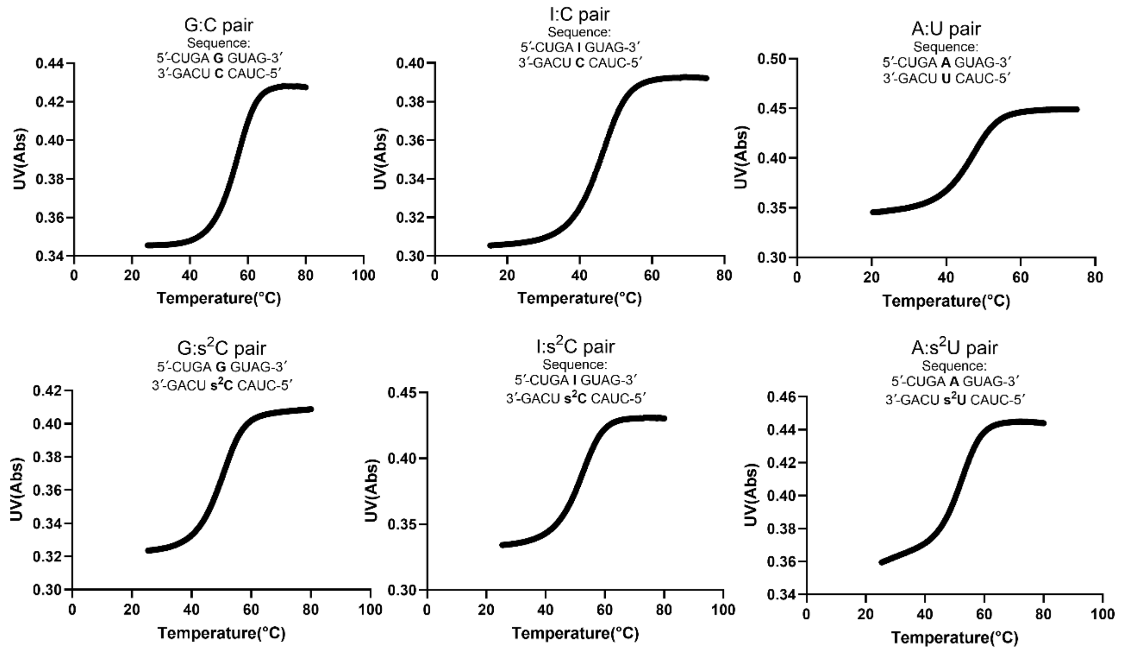


**Figure S1.** Representative melting curves collected during the thermal denaturation of 5 µM oligonucleotides in 10 mM Tris-HCl pH 8.0, 1 M NaCl, and 2.5 mM EDTA.

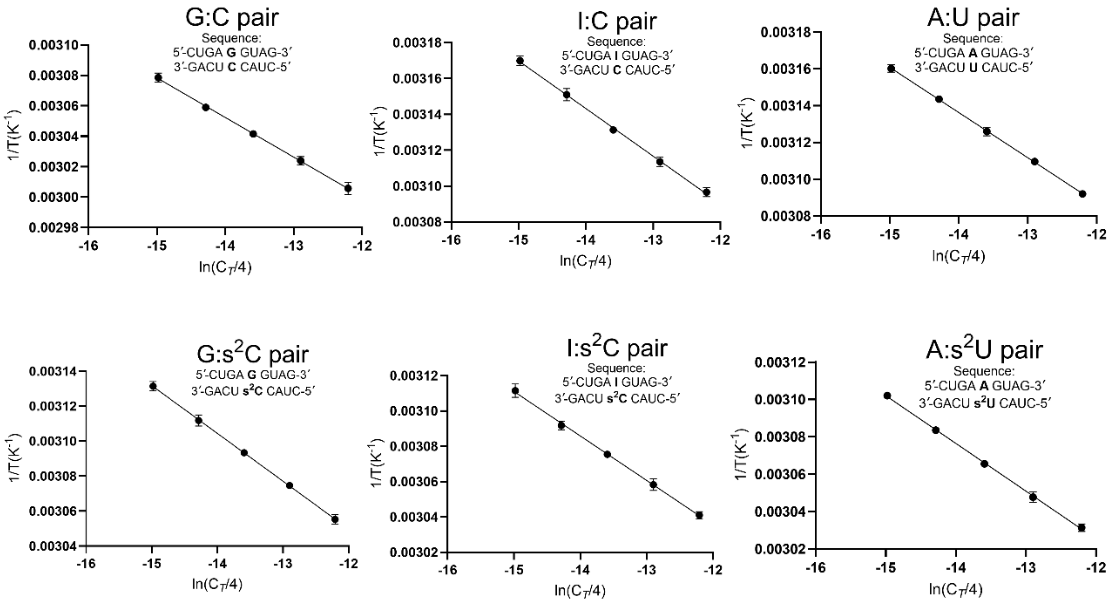


**Figure S2.** Linear least-squared fits of a Van’t Hoff plot of inverse melting temperature ($T_m^{-1}$) collected from optical melts at different oligonucleotide concentrations.

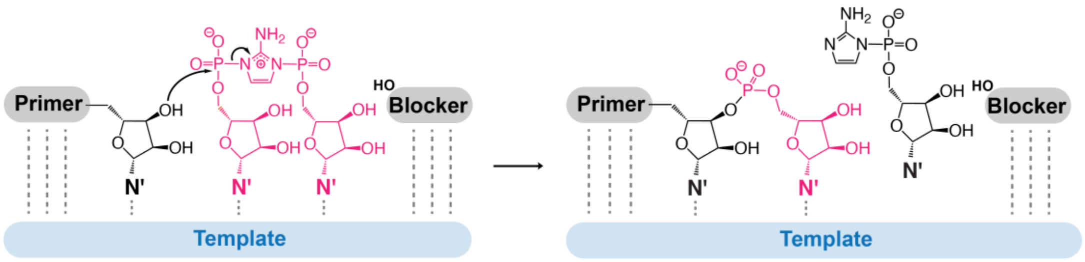


**Figure S3.** Mechanism of bridged dinucleotide (N*N) primer extension within the template-primer-blocker complex.

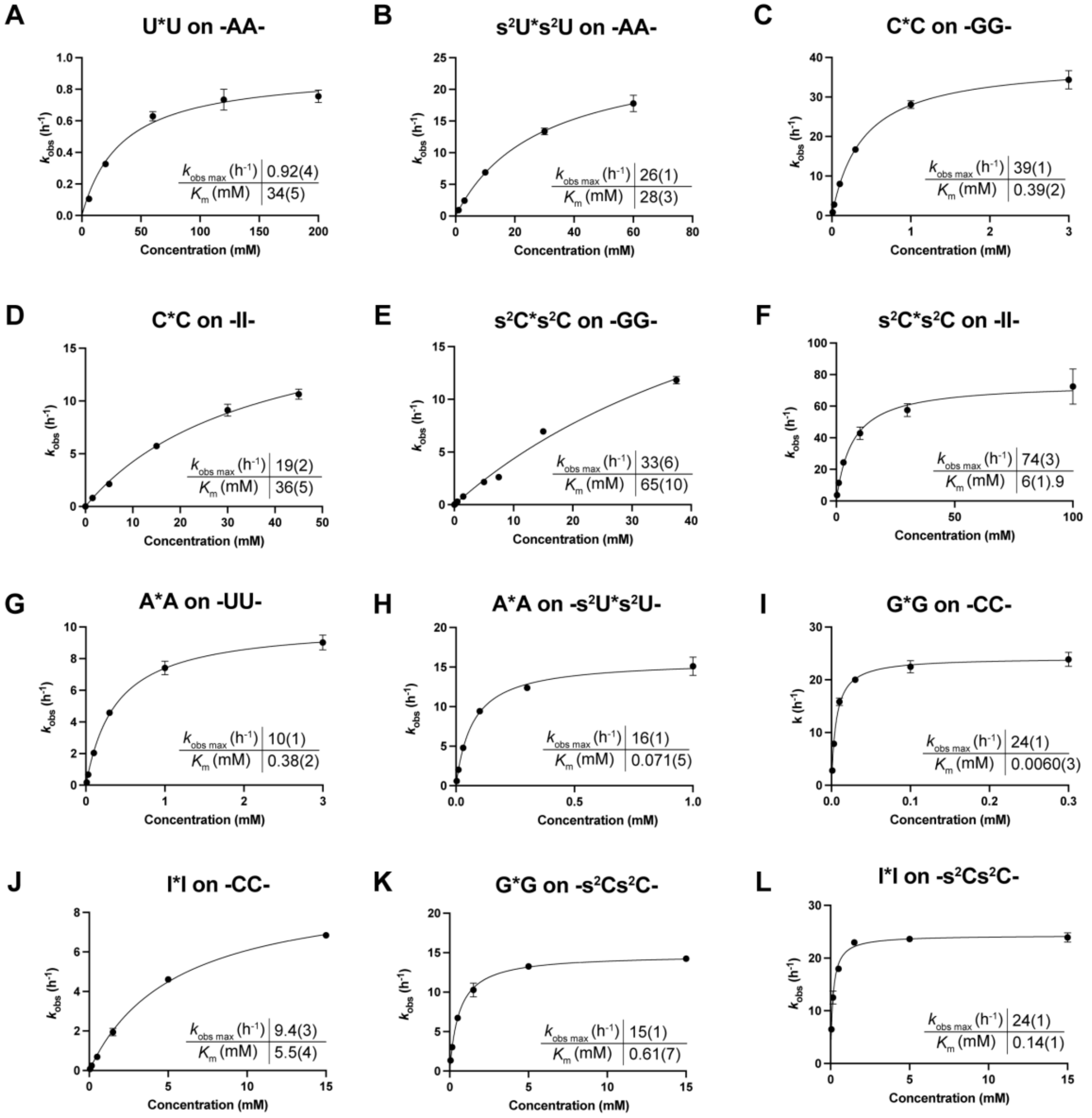


**Figure S4.** Michaelis Menten curves for primer extension reactions with bridged substrates on the indicated template sequences (A) U*U on -AA-, (B) s$^2$U*s$^2$U on -AA-, (C) C*C on -GG-, (D) C*C on -II- (E) s$^2$C*s$^2$C on -GG- (F) s$^2$C*s$^2$C on -II- (G) A*A on -UU- (H) A*A on -s$^2$Us$^2$U- (I) G*G on -CC- (J) I*I on -CC- (K) G*G on -s$^2$Cs$^2$C- (L) I*I on -s$^2$Cs$^2$C-

**Table S1. Optimized Conditions for Crystallization**

| Sequence | Optimized crystallization conditions |
|---|---|
| GCS1 | 1.4 M Sodium citrate tribasic dihydrate, 0.1 M HEPES pH 7.5 |
| GCS2 | 0.2 M Magnesium chloride hexahydrate, 0.1 M HEPES sodium pH 7.5, 30% v/v Polyethylene glycol 400 |
| ICS1 | 0.2 M Calcium acetate hydrate, 0.1 M Sodium cacodylate trihydrate pH 6.5, 18% w/v Polyethylene glycol 8,000 |
| ICS2 | 2.0 M Ammonium sulfate, 5% v/v 2-Propanol |
| Native16 | 20 % v/v Polyethylene glycol 200, 50 mM HEPES pH 7.5, 200 mM Potassium Chloride, 25 mM Magnesium Sulfate |
| AUS1 | 2.4 M Sodium malonate pH 7.0 |
| AUS2 | 0.2 M Magnesium chloride hexahydrate, 0.1 M HEPES sodium pH 7.5, 30% v/v Polyethylene glycol 400 |

**Table S2. Data Collection Statistics**

| Sequences | GCS1 | GCS2 | ICS1 | ICS2 |
|---|---|---|---|---|
| PDB code | 9CSO | 9CSP | 9CSQ | 9CSR |
| Beamline | 8.2.1 | 8.2.1 | 8.2.1 | 8.2.1 |
| Wavelength (Å) | 1.00003 | 1.00003 | 1.00003 | 1.00003 |
| Space group | *R*32 | *R*32 | *R*32 | *R*32 |
| Unit cell parameters (Å, °) | 41.34, 41.34, 124.15, 90, 90, 120 | 41.22, 41.22,123.47, 90, 90, 120 | 43.40, 43.40, 123.14, 90, 90, 120 | 40.94, 40.94, 123.48, 90, 90, 120 |
| Resolution range (Å) | 50-1.40 (1.42-1.40) | 50-1.60 (1.63-1.60) | 41.05-1.54 (1.57-1.54) | 50-1.33 (1.35-1.33) |
| Unique reflections | 8355 (389) | 5469 (228) | 6923 (339) | 9369(427) |
| Completeness (%) | 99.1 (95.6) | 97.5 (82.9) | 99.9 (100) | 98.4 (92.0) |
| $R_{merge}$ (%) | 4.8 (50.4) | 10.6 (25.4) | 5.2 (72.5) | 6.8 (47.1) |
| $\langle I/\sigma(I) \rangle$ | 23.4 (3.7) | 14.8 (2.7) | 19.0 (2.5) | 21.6 (2.5) |

| Sequences | Native16 | AUS1 | AUS2 |
|---|---|---|---|
| PDB code | 9MDW | 9MDX | 9MDY |
| Beamline | 5.0.1 | 5.0.1 | 5.0.1 |
| Wavelength (Å) | 0.97741 | 0.97741 | 0.97741 |
| Space group | *R*32 | *R*32 | *R*32 |
| Unit cell parameters (Å, °) | 41.16, 41.16, 124.20, 90, 90, 120 | 41.38, 41.38, 124.07, 90, 90, 120 | 41.00, 41.00, 122.92, 90, 90, 120 |
| Resolution range (Å) | 50-1.42 (1.44-1.42) | 50-1.50 (1.53-1.50) | 50-1.36 (1.38-1.36) |
| Unique reflections | 7975 (385) | 6663 (323) | 8888 (405) |
| Completeness (%) | 100 (100) | 96.6 (95.3) | 100 (100) |
| $R_{merge}$ (%) | 7.1 (48.1) | 5.7 (47.3) | 4.4 (49.3) |
| $\langle I/\sigma(I) \rangle$ | 24.3 (4.2) | 34.6 (4.8) | 49.4 (3.4) |

**Table S3. Data Refinement Statistics**

| Sequences | GCS1 | GCS2 | ICS1 | ICS2 |
|---|---|---|---|---|
| PDB code | 9CSO | 9CSP | 9CSQ | 9CSR |
| RNA strands per asymmetric unit | 1 | 1 | 1 | 1 |
| Resolution range (Å) | 31.0-1.40 | 34.3-1.60 | 36.0 -1.54 | 30.75-1.33 |
| Number of reflections | 8258 | 5162 | 6881 | 9192 |
| $R_{work}$ (%) | 18.9 | 17.6 | 22.2 | 18.8 |
| $R_{free}$ (%) | 22.4 | 26.4 | 24.5 | 21.8 |
| Bond length R.M.S. (Å) | 2.07 | 2.26 | 2.48 | 2.35 |
| Bond angle R.M.S. (°) | 0.020 | 0.011 | 0.013 | 0.017 |
| Average B-factors (Å$^2$) | 16.3 | 13.5 | 29.1 | 12.9 |

| Sequences | Native16 | AUS1 | AUS2 |
|---|---|---|---|
| PDB code | 9MDW | 9MDX | 9MDY |
| RNA strands per asymmetric unit | 1 | 1 | 1 |
| Resolution range (Å) | 41.4-1.42 | 34.4 -1.50 | 41.0-1.36 |
| Number of reflections | 7944 | 6647 | 8856 |
| $R_{work}$ (%) | 18.1 | 18.8 | 18.6 |
| $R_{free}$ (%) | 22.0 | 24.1 | 21.8 |
| Bond length R.M.S. (Å) | 1.98 | 0.98 | 2.25 |
| Bond angle R.M.S. (°) | 0.010 | 0.005 | 0.013 |
| Average B-factors (Å$^2$) | 11.1 | 12.9 | 12.2 |

**Table S4. Local base pair parameters for GCS1**

| Pair | Shear (Å) | Stretch (Å) | Stagger (°) | Buckle (°) | Propeller (°) | Opening (°) |
|---|---|---|---|---|---|---|
| A-U | -0.07 | -0.15 | 0.01 | -1.98 | -10.58 | -0.34 |
| G-c | -0.21 | -0.09 | -0.34 | -5.77 | -17.3 | -7.27 |
| A-U | 0.05 | -0.13 | -0.04 | -2.9 | -10.99 | 0.12 |
| G-C | -0.37 | -0.15 | -0.15 | -3.19 | -11.15 | 0.21 |
| A-U | 0.05 | -0.15 | -0.1 | -3.96 | -8.29 | 3.18 |
| A-U | -0.44 | -0.05 | -0.08 | -4.9 | -16.22 | 3.52 |
| G-C | -0.3 | -0.14 | 0.05 | -3.46 | -10.59 | 0.72 |
| A-U | -0.01 | -0.15 | -0.12 | -1.24 | -14.71 | 0.15 |
| U-A | 0.01 | -0.15 | -0.12 | 1.24 | -14.71 | 0.16 |
| C-G | 0.3 | -0.14 | 0.05 | 3.46 | -10.59 | 0.73 |
| U-A | 0.44 | -0.05 | -0.08 | 4.9 | -16.22 | 3.51 |
| U-A | -0.05 | -0.15 | -0.1 | 3.96 | -8.29 | 3.18 |
| C-G | 0.37 | -0.15 | -0.15 | 3.19 | -11.15 | 0.2 |
| U-A | -0.05 | -0.13 | -0.04 | 2.9 | -10.99 | 0.12 |
| c-G | 0.21 | -0.09 | -0.34 | 5.77 | -17.3 | -7.27 |
| U-A | 0.07 | -0.15 | 0.01 | 1.98 | -10.58 | -0.34 |

**Table S5. Local base pair step parameters for GCS1**

| Step | Shift (Å) | Slide (Å) | Rise (Å) | Tilt (°) | Roll (°) | Twist (°) | Overlap Area (Å$^2$) |
|---|---|---|---|---|---|---|---|
| AG/cU | -0.86 | -1.24 | 3.29 | -0.28 | 7.89 | 33.59 | 2.68 |
| GA/Uc | 0.88 | -1.24 | 3.13 | -0.92 | 4.44 | 33.33 | 5.51 |
| AG/CU | -0.33 | -1.42 | 3.24 | -0.59 | 10.77 | 33.07 | 1.72 |
| GA/UC | 0.38 | -1.53 | 3.27 | 0.14 | 10.48 | 30.41 | 4.78 |
| AA/UU | 0.58 | -1.81 | 3.21 | 2.58 | 14.07 | 31.51 | 2.30 |
| AG/CU | -0.15 | -2.13 | 3.07 | -0.86 | 10.5 | 25.21 | 2.30 |
| GA/UC | -0.56 | -1.53 | 3.17 | -0.11 | 5.25 | 33.48 | 3.48 |
| AU/AU | 0 | -1.15 | 3.11 | 0 | 5.56 | 32.22 | 8.33 |
| UC/GA | 0.56 | -1.53 | 3.17 | 0.11 | 5.25 | 33.48 | 3.48 |
| CU/AG | 0.15 | -2.13 | 3.07 | 0.85 | 10.5 | 25.21 | 2.30 |
| UU/AA | -0.58 | -1.81 | 3.21 | -2.58 | 14.07 | 31.5 | 2.30 |
| UC/GA | -0.38 | -1.53 | 3.27 | -0.14 | 10.48 | 30.41 | 4.78 |
| CU/AG | 0.33 | -1.42 | 3.24 | 0.59 | 10.76 | 33.08 | 1.72 |
| Uc/GA | -0.88 | -1.24 | 3.13 | 0.92 | 4.45 | 33.32 | 5.51 |
| cU/AG | 0.86 | -1.24 | 3.29 | 0.27 | 7.89 | 33.59 | 2.68 |

**Table S6. Local base pair parameters for GCS2**

| Pair | Shear (Å) | Stretch (Å) | Stagger (°) | Buckle (°) | Propeller (°) | Opening (°) |
|---|---|---|---|---|---|---|
| A-U | 0.04 | -0.19 | 0.13 | 0.55 | -10.19 | -0.24 |
| G-C | -0.28 | -0.27 | -0.07 | -2.83 | -14.2 | -0.76 |
| A-U | 0.01 | -0.21 | 0.08 | -1.97 | -11.87 | 1.79 |
| G-C | -0.49 | -0.23 | -0.22 | -2.39 | -9.86 | -1.56 |
| A-U | -0.04 | -0.16 | -0.19 | -6.94 | -10.99 | 4.53 |
| A-U | 0.13 | -0.16 | -0.25 | -10.74 | -13.63 | 1.5 |
| G-c | -0.1 | -0.06 | -0.24 | -10.72 | -12 | -8.98 |
| A-U | -0.04 | -0.21 | 0.06 | -3.1 | -12.07 | 2.05 |
| U-A | 0.04 | -0.21 | 0.06 | 3.11 | -12.08 | 2.05 |
| c-G | 0.1 | -0.06 | -0.24 | 10.72 | -12 | -8.98 |
| U-A | -0.13 | -0.16 | -0.24 | 10.73 | -13.63 | 1.49 |
| U-A | 0.04 | -0.16 | -0.19 | 6.94 | -10.99 | 4.53 |
| C-G | 0.49 | -0.23 | -0.22 | 2.39 | -9.87 | -1.57 |
| U-A | -0.01 | -0.21 | 0.08 | 1.97 | -11.87 | 1.78 |
| C-G | 0.28 | -0.27 | -0.07 | 2.83 | -14.19 | -0.76 |
| U-A | -0.04 | -0.19 | 0.13 | -0.54 | -10.19 | -0.24 |

**Table S7. Local base pair step parameters for GCS2**

| Step | Shift (Å) | Slide (Å) | Rise (Å) | Tilt (°) | Roll (°) | Twist (°) | Overlap Area (Å$^2$) |
|---|---|---|---|---|---|---|---|
| AG/CU | -0.57 | -1.18 | 3.24 | -0.84 | 6.4 | 34.16 | 2.46 |
| GA/UC | 0.64 | -1.32 | 3.15 | -0.21 | 3.48 | 33.87 | 4.97 |
| AG/CU | -0.29 | -1.63 | 3.21 | 0.46 | 11.91 | 29.17 | 1.97 |
| GA/UC | 0.57 | -1.68 | 3.36 | -0.95 | 13.31 | 31.25 | 4.42 |
| AA/UU | 0.41 | -1.69 | 3.33 | 2.86 | 12.66 | 32.83 | 2.95 |
| AG/cU | -0.64 | -1.87 | 3.23 | 1.47 | 8.96 | 27.91 | 1.73 |
| GA/Uc | 0.44 | -1.39 | 3.06 | -1.27 | 6.97 | 30.76 | 5.08 |
| AU/AU | 0 | -1.08 | 3.08 | 0 | 9.3 | 31.14 | 8.82 |
| Uc/GA | -0.44 | -1.39 | 3.06 | 1.27 | 6.97 | 30.76 | 5.08 |
| cU/AG | 0.64 | -1.87 | 3.23 | -1.47 | 8.95 | 27.9 | 1.76 |
| UU/AA | -0.41 | -1.69 | 3.33 | -2.86 | 12.66 | 32.84 | 2.95 |
| UC/GA | -0.57 | -1.68 | 3.36 | 0.95 | 13.31 | 31.24 | 4.42 |
| CU/AG | 0.29 | -1.63 | 3.21 | -0.46 | 11.91 | 29.17 | 1.97 |
| UC/GA | -0.64 | -1.32 | 3.15 | 0.21 | 3.48 | 33.87 | 4.97 |
| CU/AG | 0.57 | -1.18 | 3.24 | 0.84 | 6.4 | 34.17 | 2.46 |

**Table S8. Local base pair parameters for ICS1**

| Pair | Shear (Å) | Stretch (Å) | Stagger (°) | Buckle (°) | Propeller (°) | Opening (°) |
|---|---|---|---|---|---|---|
| A-U | 0.02 | 0.04 | -0.03 | -4.75 | -9.42 | -2.59 |
| I-c | -0.31 | -0.23 | 0.04 | 2.83 | -11.95 | -3.31 |
| A-U | 0.06 | -0.17 | 0.03 | 2.51 | -13.32 | 3.06 |
| G-C | -0.55 | -0.2 | -0.14 | -3.68 | -10.92 | 0.66 |
| A-U | -0.04 | -0.15 | -0.09 | -9.07 | -5.73 | 1.26 |
| A-U | 0.24 | -0.22 | -0.1 | -6.04 | -9.63 | -0.95 |
| G-C | -0.12 | -0.12 | 0.08 | -6.41 | -14.79 | 0.56 |
| A-U | -0.08 | -0.28 | 0.08 | -0.94 | -12.26 | 3.21 |
| U-A | 0.08 | -0.28 | 0.08 | 0.94 | -12.26 | 3.2 |
| C-G | 0.12 | -0.12 | 0.08 | 6.41 | -14.78 | 0.56 |
| U-A | -0.24 | -0.22 | -0.1 | 6.03 | -9.63 | -0.95 |
| U-A | 0.04 | -0.15 | -0.09 | 9.07 | -5.73 | 1.26 |
| C-G | 0.55 | -0.2 | -0.14 | 3.68 | -10.92 | 0.67 |
| U-A | -0.06 | -0.17 | 0.03 | -2.51 | -13.32 | 3.06 |
| c-I | 0.31 | -0.23 | 0.04 | -2.83 | -11.95 | -3.31 |
| U-A | -0.02 | 0.04 | -0.03 | 4.75 | -9.42 | -2.6 |

**Table S9. Local base pair step parameters for ICS1**

| Step | Shift (Å) | Slide (Å) | Rise (Å) | Tilt (°) | Roll (°) | Twist (°) | Overlap Area (Å$^2$) |
|---|---|---|---|---|---|---|---|
| AI/cU | -0.42 | -1.16 | 3.01 | -0.73 | 2 | 30.81 | 4.08 |
| IA/Uc | 0.62 | -1.12 | 3.23 | -0.68 | 7.79 | 34.78 | 5.12 |
| AG/CU | -0.33 | -1.69 | 3.25 | -0.97 | 12.25 | 30.82 | 1.53 |
| GA/UC | 0 | -1.74 | 3.3 | -1.69 | 13.96 | 32.14 | 3.78 |
| AA/UU | 0.16 | -1.53 | 3.19 | 2.24 | 8.9 | 32.77 | 3.17 |
| AG/CU | 0.47 | -2.01 | 3.13 | 1.56 | 9.07 | 28.29 | 3.08 |
| GA/UC | -0.28 | -1.6 | 3.09 | -0.78 | 5.6 | 30.88 | 3.62 |
| AU/AU | 0 | -1.07 | 3.1 | 0 | 8.09 | 31.78 | 8.94 |
| UC/GA | 0.28 | -1.6 | 3.09 | 0.78 | 5.6 | 30.88 | 3.62 |
| CU/AG | -0.47 | -2.01 | 3.13 | -1.57 | 9.07 | 28.28 | 3.08 |
| UU/AA | -0.16 | -1.53 | 3.19 | -2.23 | 8.9 | 32.76 | 3.17 |
| UC/GA | 0 | -1.74 | 3.3 | 1.69 | 13.96 | 32.14 | 3.78 |
| CU/AG | 0.33 | -1.69 | 3.25 | 0.97 | 12.25 | 30.82 | 1.53 |
| Uc/IA | -0.62 | -1.12 | 3.23 | 0.68 | 7.79 | 34.78 | 5.12 |
| cU/AI | 0.42 | -1.16 | 3.02 | 0.72 | 1.99 | 30.8 | 4.08 |

**Table S10. Local base pair parameters for ICS2**

| Pair | Shear (Å) | Stretch (Å) | Stagger (°) | Buckle (°) | Propeller (°) | Opening (°) |
|---|---|---|---|---|---|---|
| A-U | -0.07 | -0.19 | 0.07 | -0.78 | -10.33 | 0.7 |
| G-C | -0.23 | -0.26 | -0.05 | -3.29 | -13.84 | -1.61 |
| A-U | 0.07 | -0.16 | 0.02 | -2.32 | -13.34 | 2.46 |
| G-C | -0.34 | -0.22 | -0.07 | -1.24 | -11.34 | -1.55 |
| A-U | 0.05 | -0.21 | -0.1 | -4.04 | -8.17 | 2.39 |
| A-U | -0.03 | -0.03 | -0.09 | -11.06 | -16.06 | 3.75 |
| I-c | -0.28 | -0.2 | 0.04 | -1.61 | -8.43 | 0.93 |
| A-U | -0.05 | -0.21 | 0.02 | 0.24 | -17.28 | 1.38 |
| U-A | 0.05 | -0.21 | 0.02 | -0.24 | -17.28 | 1.38 |
| c-I | 0.28 | -0.2 | 0.04 | 1.61 | -8.43 | 0.93 |
| U-A | 0.03 | -0.03 | -0.09 | 11.06 | -16.06 | 3.75 |
| U-A | -0.05 | -0.21 | -0.1 | 4.04 | -8.17 | 2.39 |
| C-G | 0.34 | -0.22 | -0.07 | 1.24 | -11.34 | -1.55 |
| U-A | -0.07 | -0.16 | 0.02 | 2.32 | -13.34 | 2.46 |
| C-G | 0.23 | -0.26 | -0.05 | 3.29 | -13.84 | -1.61 |
| U-A | 0.07 | -0.19 | 0.07 | 0.78 | -10.33 | 0.7 |

**Table S11. Local base pair step parameters for ICS2**

| Step | Shift (Å) | Slide (Å) | Rise (Å) | Tilt (°) | Roll (°) | Twist (°) | Overlap Area (Å$^2$) |
|---|---|---|---|---|---|---|---|
| AG/CU | -0.6 | -1.07 | 3.23 | -1.08 | 6.88 | 33.94 | 3.13 |
| GA/UC | 0.69 | -1.2 | 3.18 | 0.67 | 5.02 | 33.98 | 5.23 |
| AG/CU | -0.52 | -1.4 | 3.16 | -1.82 | 10.96 | 31.88 | 1.90 |
| GA/UC | 0.47 | -1.61 | 3.31 | 0.26 | 11.14 | 29.83 | 4.71 |
| AA/UU | 0.87 | -1.79 | 3.36 | 3.85 | 15.78 | 31.87 | 3.51 |
| AI/cU | -0.64 | -1.79 | 2.92 | -1.02 | 7.46 | 25.91 | 1.97 |
| IA/Uc | -0.13 | -1.33 | 3.2 | 0.03 | 8.03 | 32.79 | 3.32 |
| AU/AU | 0 | -1.24 | 3.2 | 0 | 6.34 | 31.7 | 8.83 |
| Uc/IA | 0.13 | -1.33 | 3.2 | -0.03 | 8.03 | 32.79 | 3.32 |
| cU/AI | 0.64 | -1.79 | 2.92 | 1.02 | 7.46 | 25.91 | 1.97 |
| UU/AA | -0.87 | -1.79 | 3.36 | -3.85 | 15.78 | 31.87 | 3.51 |
| UC/GA | -0.47 | -1.61 | 3.31 | -0.26 | 11.14 | 29.83 | 4.71 |
| CU/AG | 0.52 | -1.4 | 3.16 | 1.82 | 10.96 | 31.88 | 1.90 |
| UC/GA | -0.69 | -1.2 | 3.18 | -0.67 | 5.02 | 33.98 | 5.23 |
| CU/AG | 0.6 | -1.07 | 3.23 | 1.08 | 6.88 | 33.94 | 3.13 |

**Table S12. Local base pair parameters for native sequence Native16**

| Pair | Shear (Å) | Stretch (Å) | Stagger (°) | Buckle (°) | Propeller (°) | Opening (°) |
|---|---|---|---|---|---|---|
| A-U | -0.07 | -0.12 | 0.05 | 0.18 | -9.04 | 0.61 |
| G-C | -0.21 | -0.22 | -0.13 | -4.46 | -14.88 | -1.3 |
| A-U | 0.03 | -0.09 | 0.03 | -3.73 | -12.03 | 0.75 |
| G-C | -0.41 | -0.18 | -0.14 | -2.66 | -10.98 | -1.84 |
| A-U | 0.07 | -0.21 | -0.11 | -6.42 | -8.83 | 1.77 |
| A-U | 0.12 | -0.06 | -0.18 | -9.07 | -14.32 | 1.15 |
| G-C | -0.15 | -0.15 | 0.06 | -4.43 | -10.57 | -0.33 |
| A-U | -0.02 | -0.13 | -0.02 | -0.35 | -12.71 | 0.28 |
| U-A | 0.02 | -0.13 | -0.02 | 0.35 | -12.71 | 0.29 |
| C-G | 0.15 | -0.15 | 0.06 | 4.43 | -10.57 | -0.33 |
| U-A | -0.12 | -0.06 | -0.18 | 9.08 | -14.31 | 1.15 |
| U-A | -0.07 | -0.21 | -0.11 | 6.41 | -8.83 | 1.77 |
| C-G | 0.41 | -0.18 | -0.14 | 2.66 | -10.98 | -1.85 |
| U-A | -0.03 | -0.09 | 0.03 | 3.74 | -12.03 | 0.75 |
| C-G | 0.21 | -0.22 | -0.13 | 4.46 | -14.88 | -1.3 |
| U-A | 0.07 | -0.12 | 0.05 | -0.18 | -9.05 | 0.6 |

**Table S13. Local base pair step parameters for native sequence Native16**

| Step | Shift (Å) | Slide (Å) | Rise (Å) | Tilt (°) | Roll (°) | Twist (°) | Overlap Area ($Å^2$) |
|---|---|---|---|---|---|---|---|
| AG/CU | -0.64 | -1.16 | 3.3 | -0.6 | 7.58 | 34.43 | 2.72 |
| GA/UC | 0.58 | -1.2 | 3.17 | -0.09 | 4.29 | 33.88 | 5.22 |
| AG/CU | -0.24 | -1.5 | 3.2 | -0.25 | 10.77 | 30.82 | 2.13 |
| GA/UC | 0.53 | -1.65 | 3.38 | 0.07 | 11.31 | 31.19 | 4.57 |
| AA/UU | 0.68 | -1.62 | 3.34 | 4.46 | 14.5 | 31.98 | 3.73 |
| AG/CU | -0.03 | -2.22 | 3.05 | -0.39 | 10.57 | 24.81 | 2.38 |
| GA/UC | -0.55 | -1.4 | 3.15 | -0.48 | 4.59 | 33.3 | 3.72 |
| AU/AU | 0 | -1.06 | 3.2 | 0 | 9.96 | 31.64 | 8.54 |
| UC/GA | 0.55 | -1.4 | 3.15 | 0.48 | 4.59 | 33.3 | 3.72 |
| CU/AG | 0.03 | -2.22 | 3.05 | 0.39 | 10.58 | 24.81 | 2.38 |
| UU/AA | -0.68 | -1.62 | 3.34 | -4.46 | 14.5 | 31.98 | 3.73 |
| UC/GA | -0.53 | -1.65 | 3.38 | -0.07 | 11.31 | 31.19 | 4.57 |
| CU/AG | 0.24 | -1.5 | 3.2 | 0.25 | 10.77 | 30.81 | 2.13 |
| UC/GA | -0.58 | -1.2 | 3.18 | 0.09 | 4.29 | 33.88 | 5.22 |
| CU/AG | 0.64 | -1.16 | 3.3 | 0.6 | 7.58 | 34.43 | 2.72 |

**Table S14. Combination of the primer, template, blocker and complementary oligonucleotides used in the Michaelis-Menten analysis of primer extension reactions.**

| Bridged dinucleotide | Template Sequence | Primer | Template | Blocker | Complementary DNA |
|---|---|---|---|---|---|
| U*U | -AA- | DL-30 | LA-121 | LA-111 | dCLA-121 |
| $s^2U*s^2U$ | -AA- | DL-30 | LA-121 | LA-111 | dCLA-121 |
| C*C | -GG- | DL-30 | LA-124 | LA-111 | dCLA-123 |
| C*C | -II- | DL-30 | IIT26 | LA-111 | dCLA-123 |
| $s^2C*s^2C$ | -GG- | DL-30 | LA-124 | LA-111 | dCLA-124 |
| $s^2C*s^2C$ | -II- | DL-30 | IIT26 | LA-111 | dCLA-123 |
| A*A | -UU- | DL-30 | LA-122 | LA-111 | dCLA-122 |
| A*A | $-s^2Us^2U-$ | DL-30 | S2UT | LA-111 | dCLA-122 |
| G*G | -CC- | DL-30 | LA-123 | LA-111 | dCLA-124 |
| I*I | -CC- | DL-30 | LA-123 | LA-111 | dCLA-123 |
| G*G | $-s^2Cs^2C-$ | DL-30 | S2CT | LA-111 | dCLA-123 |
| I*I | $-s^2Cs^2C-$ | DL-30 | S2CT | LA-111 | dCLA-123 |

**Table S15. Sequences of oligonucleotides used in the Michaelis-Menten analysis of primer extension reactions.**

| Name | Role | Source | Type | Sequence (5'→ 3') |
|---|---|---|---|---|
| DL-30 | Primer | IDT | RNA | /FAM/AGU GAG UAA CGG |
| LA-111 | Blocker | IDT | RNA | G AUG UCA GAU AU |
| IIT26 | Template | In-house | RNA | AU AUC UGA CAU C**II** CCG UUA CUC ACU |
| LA-121 | Template | IDT | RNA | AU AUC UGA CAU C**AA** CCG UUA CUC ACU |
| dCLA-121 | Complementary strand | IDT | DNA | AGT GAG TAA CGG **TT**G ATG TCA GAT AT |
| LA-122 | Template | IDT | RNA | AU AUC UGA CAU C**UU** CCG UUA CUC ACU |
| dCLA-122 | Complementary strand | IDT | DNA | AGT GAG TAA CGG **AA**G ATG TCA GAT AT |
| LA-123 | Template | IDT | RNA | AU AUC UGA CAU C**CC** CCG UUA CUC ACU |
| dCLA-123 | Complementary strand | IDT | DNA | AGT GAG TAA CGG **GG**G ATG TCA GAT AT |
| LA-124 | Template | IDT | RNA | AU AUC UGA CAU C**GG** CCG UUA CUC ACU |
| dCLA-124 | Complementary strand | IDT | DNA | AGT GAG TAA CGG **CC**G ATG TCA GAT AT |
| 2SCT | Template | In-house | RNA | AU AUC UGA CAU C**s$^2$Cs$^2$C** CCG UUA CUC ACU |
| 2SUT | Template | In-house | RNA | AU AUC UGA CAU C**s$^2$Us$^2$U** CCG UUA CUC ACU |

### References


1. D. Ding, L. Zhou, C. Giurgiu, J. W. Szostak, Kinetic explanations for the sequence biases observed in the nonenzymatic copying of RNA templates. *Nucleic Acids Research* **50**, 35-45 (2022).
